# The transcription factor Bhlhe40 promotes inflammatory functions of ILC2s through inducing GM-CSF while inhibiting IL-10 expression

**DOI:** 10.64898/2026.08.23.746557

**Authors:** Xiaoliang Zhu, Jian Song, Jia Nie, Xi Chen, Yaqiang Cao, Danping Wei, Rama K. Gurram, Dingkang Peng, Keji Zhao, Remy Bosselut, Jinfang Zhu

## Abstract

Group 2 innate lymphoid cells (ILC2s) regulate type 2 immune responses partly by recruiting eosinophils, but the mechanisms underlying this process remain incompletely understood. Although both ILC2s and type 2 T helper (Th2) cells are capable of expressing IL-5, a cytokine critical for recruiting eosinophils, ILC2s are more potent than Th2 cells in this process. Here, we show that the transcription factor Bhlhe40 promotes GM-CSF production in ILC2s, and Bhlhe40 expression in ILC2s is required for efficient eosinophil recruitment during both papain-and helminth-induced type 2 immune responses. However, Bhlhe40 deficiency in ILC2s had no effect on the production of classical type 2 cytokines, including IL-4, IL-5 and IL-13, despite Bhlhe40 being required for type 2 cytokine production by Th2 cells. Furthermore, in contrast to ILC2s, Th2 cells produced little GM-CSF and administration of GM-CSF rescued eosinophil recruitment in ILC2-specific Bhlhe40-deficient mice. In addition to promoting GM-CSF expression, Bhlhe40 repressed IL-10 production in ILC2s as it did in Th2 cells, particularly during chronic inflammation. Single-cell transcriptomic analyses further supported this regulatory network, and the ChIP-Seq data revealed direct binding of Bhlhe40 to the enhancer and promoter regions within the *Il10* and *Csf2* loci. Collectively, our findings show that Bhlhe40 exerts distinct gene regulatory functions in ILC2s and Th2 cells, and Bhlhe40 modulates the inflammatory and anti-inflammatory functions of ILC2s by promoting GM-CSF while suppressing IL-10 production.

---

Two decades ago, a non-B/Non-T cell population capable of producing type 2 cytokines was reported ^1^. Soon after that, four groups independently characterized this unique population of immune cells, now known as group 2 innate lymphoid cells (ILC2s)^2, 3, 4, 5^. Since then, ILC2s have been known to play an important role during type 2 immune responses including host defense against helminth infection, allergic inflammation, and tissue repair. Unlike adaptive type 2 T helper (Th2) cells, ILC2s lack antigen-specific receptors. However, they can respond rapidly to tissue-derived alarmins such as interleukin (IL)-25, IL-33, and thymic stromal lymphopoietin (TSLP) secreted by epithelial cells after infection or tissue injury ^6^. Upon activation, ILC2s produce large amounts of type 2 cytokines including IL-5 and IL-13, which promote eosinophil recruitment, goblet cell hyperplasia, mucus production, and airway hyperresponsiveness ^7^. The development and maintenance of ILC2 lineage identity depend on key transcription factors including GATA3, RORα, BCL11B and ID2, which collectively regulate ILC2 development and effector functions ^8, 9, 10, 11, 12^. Increasing evidence indicates that ILC2s not only initiate early innate responses but also shape adaptive immunity and chronic inflammation.

Among the cytokines produced by ILC2s, IL-5 has been recognized as a major mediator of eosinophil recruitment and survival during type 2 immune responses ^13^. Granulocyte-macrophage colony-stimulating factor (GM-CSF) is a pleiotropic cytokine originally characterized as a hematopoietic growth factor that promotes granulocyte and macrophage differentiation but now is also recognized as a central mediator for tissue inflammation by regulating the activation and survival of multiple immune cell populations ^14, 15^. ILC2s were recently reported producing GM-CSF during allergic airway inflammation and in salivary glands ^16, 17^, but its precise role during type 2 inflammation remains incompletely understood. Particularly, whether GM-CSF contributes to eosinophilic inflammation together with IL-5 has not been fully clarified. Furthermore, the transcriptional mechanisms controlling GM-CSF production in ILC2s remain unknown.

In addition to pro-inflammatory cytokines, ILC2s are also capable of producing the anti-inflammatory cytokine IL-10 under certain inflammatory conditions ^18^. IL-10-producing ILC2s have been proposed to represent a regulatory subset that restrains excessive type 2 inflammation and promotes resolution of tissue damage. Thus, the balance between the production of inflammatory and anti-inflammatory cytokines by ILC2s is likely to be an important event in regulating chronic allergic responses. While several environmental signals, including retinoic acid and IL-2, have been implicated in promoting IL-10 production in ILC2s, the intrinsic transcriptional regulators governing IL-10 expression in these cells are not well defined ^18^. Understanding how IL-10 is regulated in ILC2s may provide an important insight into mechanisms controlling pathogenic versus regulatory properties of ILC2s in type 2 immunity.

Basic helix-loop-helix family member e40 (Bhlhe40) has emerged as an important transcription factor in regulating immune cell activation and cytokine production. Bhlhe40 is broadly expressed by multiple immune cell populations, including T cells, B cells and macrophages ^19^. In CD4 T cells, Bhlhe40 promotes inflammatory cytokine production and suppresses IL-10 expression, thereby supporting pathogenic effector responses during infection and autoimmune inflammation ^20, 21, 22^. It has also been reported that Bhlhe40 inhibits germinal center response by repressing B cells and follicular helper T cells ^23, 24^. In macrophages and CD8 T cells, Bhlhe40 has been implicated in regulating inflammatory activation and cellular metabolism ^25, 26^. Recently, it has been reported that Bhlhe40 regulates effector cytokine programs in group 3 innate lymphoid cells ^27^. Despite these established functions of Bhlhe40, its role in ILC2 biology remains unknown.

ILC2s and Th2 cells function cooperatively during type 2 immune responses and engage in extensive bidirectional crosstalk. ILC2s can promote Th2 cell differentiation through IL-13 production and antigen presentation, whereas Th2-derived cytokines including IL-2 and IL-4 support ILC2 expansion and survival ^28, 29, 30^. Given the established role of Bhlhe40 in regulating cytokine production in T cells, an important unresolved question is whether Bhlhe40 controls inflammatory programs in ILC2s and Th2 cells through similar mechanisms. Defining the role of Bhlhe40 in ILC2-mediated immunity may therefore provide new insights into the transcriptional regulation of type 2 inflammation and identify potential therapeutic targets for allergic disease.

In this study, we report that Bhlhe40 represses IL-10 but promotes GM-CSF in ILC2s and Bhlhe40 expression in ILC2s is critical for efficient eosinophils recruitment during type 2 inflammation. These observations were initially identified from studying the *Bhlhe40* germline knockout mice and subsequently confirmed in an ILC2-specific *Bhlhe40* conditional knockout mouse model. Furthermore, we discovered that Bhlhe40 regulates eosinophils recruitment through differential regulation of cytokine production in Th2 cells and ILC2s. In Th2 cells, Bhlhe40 promotes the production of IL-5 and IL-13. In contrast, Bhlhe40 has no effect on production of these type 2 cytokines in ILC2s but instead promotes GM-CSF expression. Nevertheless, suppression of IL-10 production is a common regulatory function of Bhlhe40 in both ILC2s and Th2 cells. Administration of GM-CSF rescued the defect in eosinophil recruitment caused by deficiency of Bhlhe40 in ILC2s indicating that GM-CSF and IL-5 produced by ILC2s collaboratively regulate eosinophilia.

## RESULTS

### Bhlhe40 expression by ILC2s during type 2 immune responses

Activated T cells including Th2 cells express Bhlhe40 ^24^ and Bhlhe40 had been reported to regulate the expression of type 2 cytokines including IL-5 and IL-13 in Th2 cells ^31^. Since ILC2s, which also secrete type 2 cytokines, are the innate counterpart of Th2 cells, we first investigated whether ILC2s express Bhlhe40 by using the Bhlhe40-V5 mice and anti-V5 staining as we previously reported ^24^. Two major classical mouse models for type 2 immune responses as shown in **Fig. 1a** and **Fig. 1b** were used in this study: one is helminth infection and the other is papain-induced lung inflammation. Mice acutely challenged with papain for three days were analyzed for innate type 2 immune response, whereas mice chronically challenged with papain for sixteen days were used to assess both innate and adaptive type 2 immune response (**Fig. 1b**). In steady state, a large portion of ILC2s from small intestine expressed Bhlhe40 (45.97%), while lung ILC2s barely expressed Bhlhe40 (**Fig. 1c** and **Fig. 1f**). On the other hand, in mice infected with *Nippostrongylus brasiliensis* (*Nb*) for 10 to 12 days as shown in **Fig. 1a**, vast majority (83.26%) of lung ILC2s expressed Bhlhe40 (**Fig. 1d** and **Fig. 1g**). In this model, Bhlhe40 expression was also detected in Th2 cells from lung (37.56%) and from small intestine (75.35%). In mice chronically challenged with papain as shown in **Fig. 1b**, ILC2s from both lung (33.73%) and bronchoalveolar lavage fluid (BALF) (26.01%) expressed Bhlhe40, similar to Bhlhe40 expression in Th2 cells (**Fig. 1e** and **Fig. 1h**). Bhlhe40 was also expressed in ILC1s (**Supplementary Fig. 1a**) and ILC3s (**Supplementary Fig. 1b**) from small intestine in steady state. In summary, resting ILC2s, particularly from the lung tissue in steady state, express little Bhlhe40, and Bhlhe40 expression is induced after ILC2 activation by either *Nb* infection or papain treatment; this mirrors the regulation of Bhlhe40 expression during T cell activation as previously reported ^24^.

**Fig. 1.**
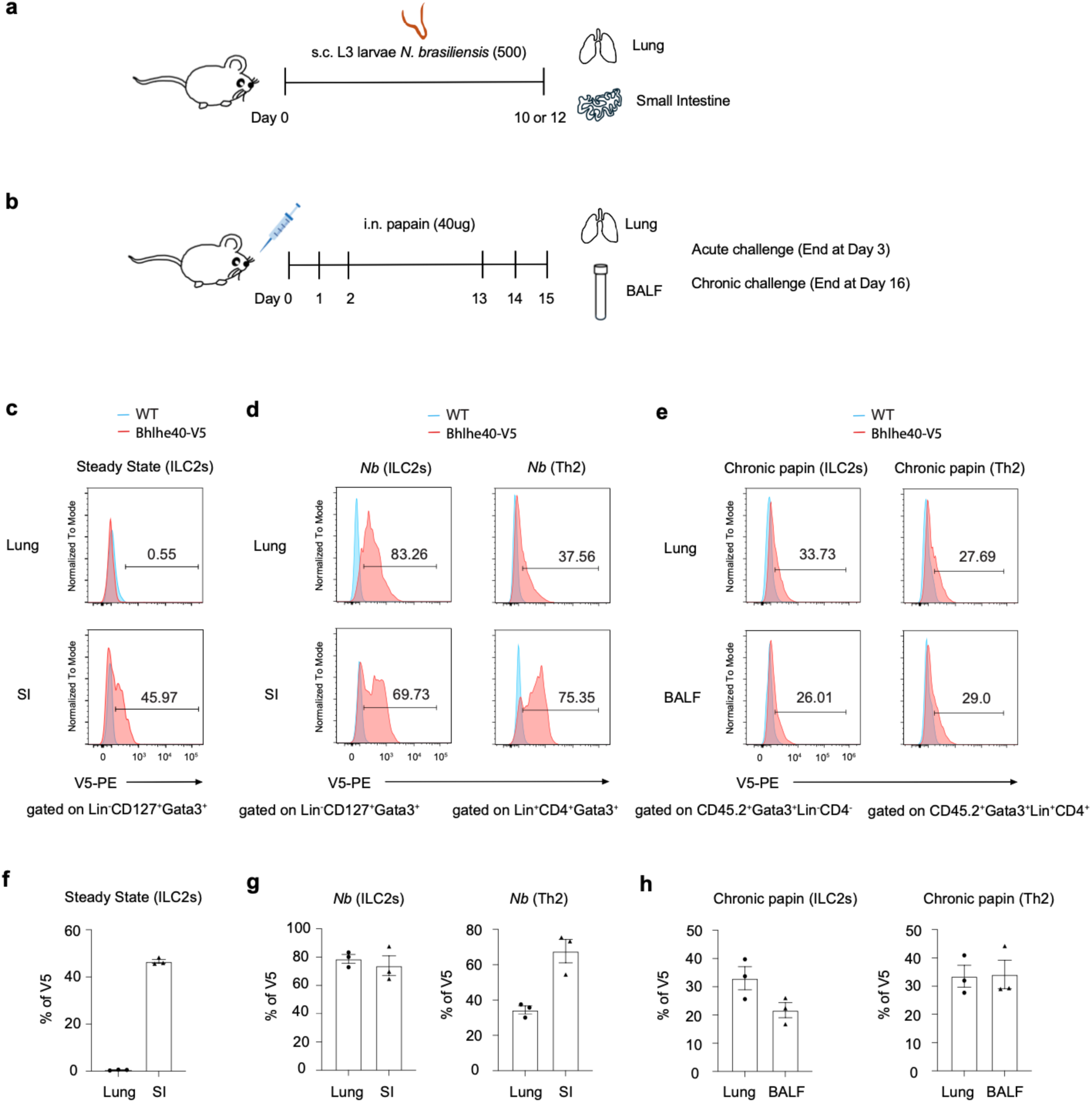
Activated ILC2s express Bhlhe40. **a**, Experimental procedure for mouse model of *N. brasiliensis* infection. **b**, Experimental procedure for acute and chronic papain challenging models. **c**, Bhlhe40 expression in ILC2s (Lin^-^CD127^+^GATA3^+^) was assessed by anti-V5 staining in ILC2s from lung and small intestine (SI) of Bhlhe40-V5 mice in steady state. **d**, WT and Bhlhe40-V5 mice were infected with 500 L3 *N. brasiliensis* (s.c.) for 10-12 days, and Bhlhe40 expression in ILC2s (Lin^-^CD127^+^GATA3^+^) and Th2 (Lin^+^CD4^+^GATA3^+^) were shown by anti-V5 staining in lung and SI. **e**, WT and Bhlhe40-V5 mice were chronically challenged with papain for 16 days, and Bhlhe40 expression in ILC2s (CD45.2^+^ GATA3^+^ Lin^-^CD4^-^) and Th2 (CD45.2^+^ GATA3^+^ Lin^+^CD4^+^) were shown by anti-V5 staining in lung and bronchoalveolar lavage fluid (BALF). **f**, Summary of percentage of V5-expressed ILC2s in **c**. **g**, Summary of percentage of V5-expressed ILC2s and Th2 in **d**. **h**, Summary of percentage of V5-expressed ILC2s and Th2 in **e**. **c**-**h** show summarized results from two independent experiments with Bhlhe40-V5 (*n* = 3) mice.

### Defective anti-helminth responses in the Bhlhe40-deficient mice

To investigate the function of Bhlhe40 in vivo during helminth infection, we infected WT and Bhlhe40 germline knockout mice (*Bhlhe40*^-/-^) with *Nb* as shown in **Fig. 2a**, then collected small intestine 10 days post infection to count adult worms. Compared with WT mice, Bhlhe40-deficient mice showed a higher worm burden (**Fig. 2a**). Eosinophils were dramatically reduced in lung, BALF, and mesenteric lymph nodes (mLNs) with Bhlhe40 deficiency (**Fig. 2b-2c**). However, both percentage and total cell number of ILC2s in Bhlhe40-deficient mice appeared to be normal (**Fig. 2d**). As shown in **Fig. 2e**, Th2 cells were found reduced in lung and BALF, but increased in mLNs, a similar phenotype that was found in ILC2-deficient mice ^28^. These data indicate that Bhlhe40 may affect the functions of ILC2s rather than ILC2 development and expansion.

**Fig. 2.**
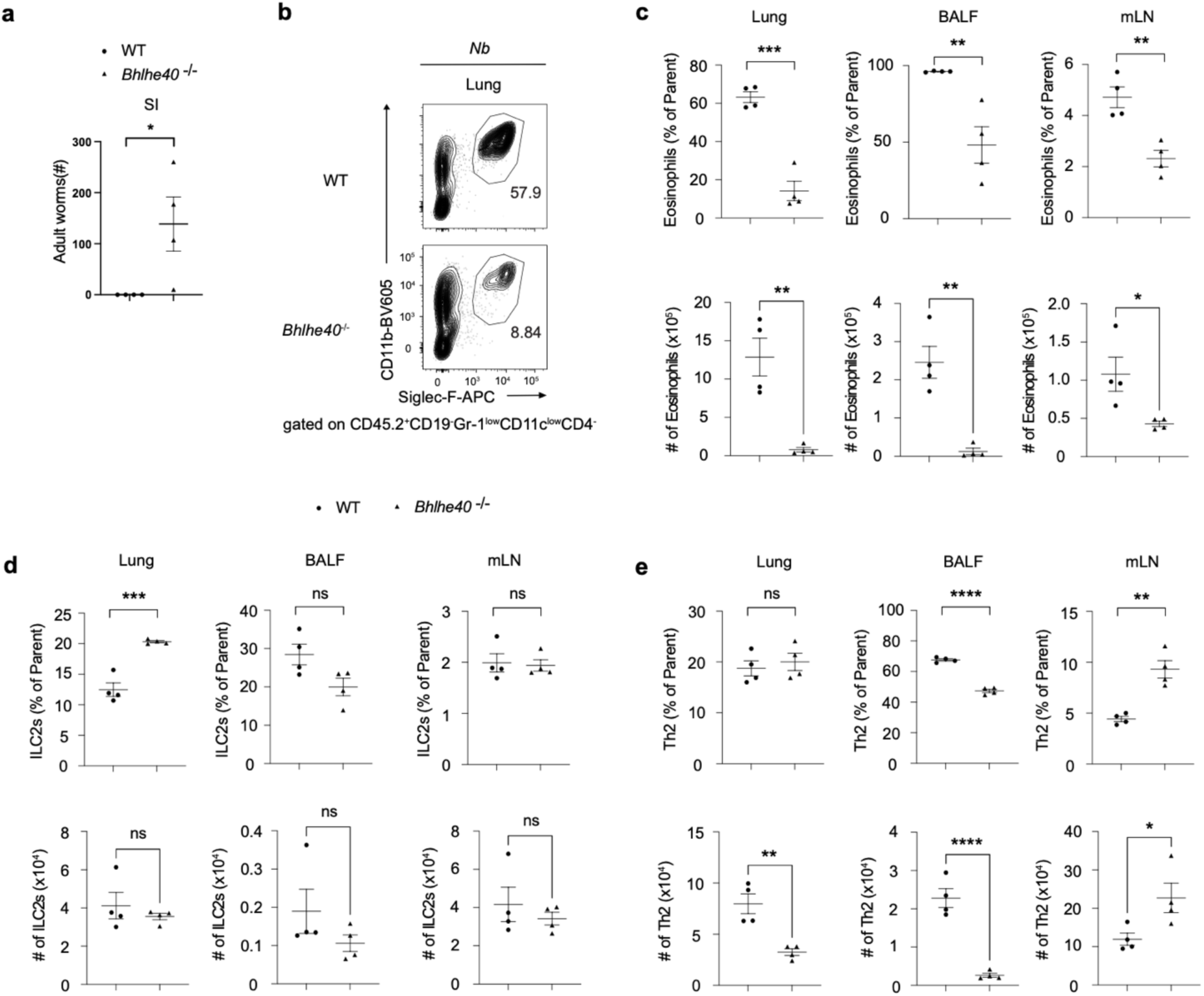
Bhlhe40 deficiency increases worm burden in host infected with *N. brasiliensis*. **a**-**e**, WT and *Bhlhe40*^-/-^ mice were infected with 500 L3 *N. brasiliensis* (s.c.) for 10 days. **a**, Adult worms in small intestine were counted. **b**, Eosinophils (CD45.2^+^CD19^-^Gr-1^low^CD11c^low^CD4^-^Siglec-F^+^CD11b^+^) in lung were analyzed by flow cytometry. **c**, Percentage and cell number of eosinophils in lung, BALF and mesenteric lymph nodes (mLNs) were summarized. **d**, Percentage and cell number of ILC2s in lung, BALF and mLNs were summarized. **e**, Percentage and cell number of Th2 in lung, BALF and mLNs were summarized. **a**-**e** show summarized results from two independent experiments with WT (*n* = 4) and *Bhlhe40*^-/-^ (*n* = 4) mice. * p < 0.05, ** p < 0.01, *** p < 0.001, **** p < 0.0001, Student’s *t*-test. Error bars indicate SEM.

To further confirm that Bhlhe40 is required for eosinophils recruitment during type 2 immune response, we challenged WT and *Bhlhe40*^-/-^ mice with chronic papain treatment and found that eosinophils were also reduced in the absence of Bhlhe40 in this model (**Supplementary Fig. 2a**), while percentages of ILC2s and Th2 cells showed no difference.

### Bhlhe40 deficiency in ILC2s reduces papain-induced type 2 immune responses

Bhlhe40 was reported to have functions in multiple types of immune cells, including CD4 T cells, T cells, B cells and macrophages ^19^. To exclude potential functions of Bhlhe40 in these cells contributing to eosinophil recruitment, we generated ILC2-specific *Bhlhe40* knockout mouse model (*Bhlhe40*^fl/fl^-*Klrg1*-Cre) by crossing *Bhlhe40*-flox mice with *Klrg1*-Cre mice, which allow gene deletion specifically in ILC2s but not in Th2 cells ^28^. We first tested the functions of Bhlhe40 in ILC2s by using the mouse model of acute papain treatment for 3 days as shown in **Fig. 1b** and found that eosinophils were reduced in both lung (**Fig. 3a** and **Fig. 3b**) and BALF (**Supplementary Fig. 2b**) from the ILC2-specific *Bhlhe40* conditional knockout mice compared with WT mice. By contrast, the percentage or total cell number of ILC2s in lung and BALF were not changed (**Fig. 3c**). These data indicate that Bhlhe40 expression in ILC2s is critical for optimal eosinophils recruitment during type 2 immune responses.

**Fig. 3.**
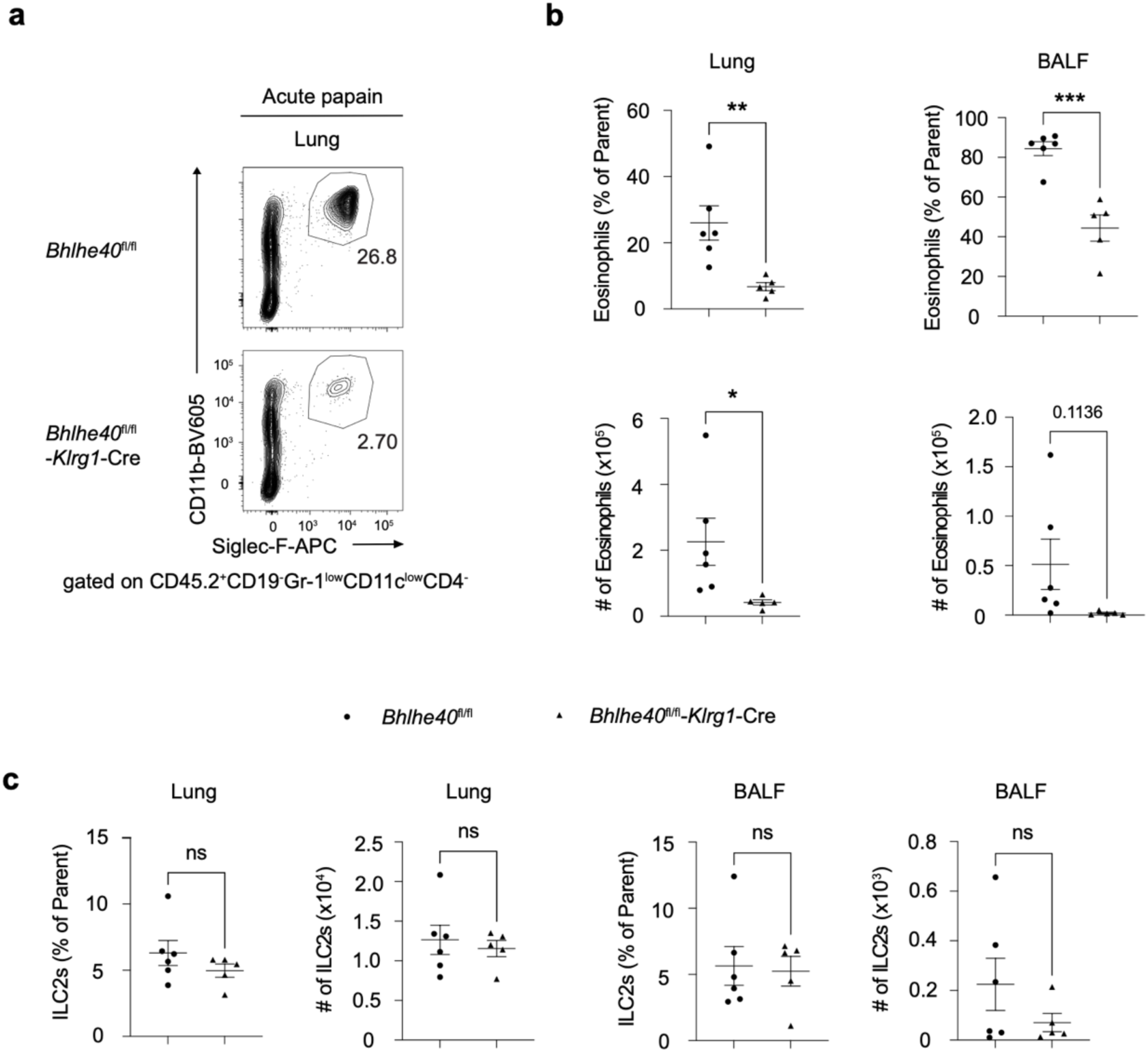
Bhlhe40 deficiency in ILC2s reduces papain-induced recruitment of eosinophils. **a**-**c**, *Bhlhe40*^fl/fl^ and *Bhlhe40*^fl/fl^-*Klrg1*-Cre mice were acutely challenged with papain for 3 days. **a**, Eosinophils (CD45.2^+^CD19^-^Gr-1^low^CD11c^low^CD4^-^Siglec-F^+^CD11b^+^) in lung were analyzed by flow cytometry. **b**, Percentage and cell number of eosinophils in lung and BALF were summarized. **c**, Percentage and cell number of ILC2s in lung and BALF were summarized. **a**-**c** show summarized results from two independent experiments with *Bhlhe40*^fl/fl^ (*n* = 6) and *Bhlhe40*^fl/fl^-*Klrg1*-Cre (*n* = 5) mice. * p < 0.05, ** p < 0.01, *** p < 0.001, **** p < 0.0001, Student’s *t*-test. Error bars indicate SEM.

### Bhlhe40 induces GM-CSF but represses IL-10 in ILC2s

Since Bhlhe40 deficiency in ILC2s has no effect on ILC2 maintenance and expansion, we turned to evaluate Bhlhe40’s effect on ILC2’s functions by testing cytokine production. The gating strategies for eosinophils and cytokines in ILC2s and Th2 were shown in **Supplementary Fig. 3a** and **Supplementary Fig. 3b**. As shown in **Fig. 4a** and **Supplementary Fig. 4a** in the steady state, very few eosinophils were found in lung, and there was no difference in eosinophils between *Bhlhe40*^fl/fl^ and *Bhlhe40*^fl/fl^-*Klrg1*-Cre mice. ILC2s were in these mice were also similar (**Supplementary Fig. 4b**). However, compared with *Bhlhe40*^fl/fl^ ILC2s, *Bhlhe40*^fl/fl^-*Klrg1*-Cre ILC2s secreted significant less GM-CSF in lung, while the production of other cytokines including IL-5, IL-13 and IL-4 was not significantly different (**Fig. 4a** and **Supplementary Fig. 4c-4d**). Very little IL-10 production was found in ILC2s in steady state. In mice challenged with papain for 3 days, much less eosinophils were recruited to the lung after *Bhlhe40* deletion in ILC2s (**Fig. 4b** and **Supplementary Fig. 4e**), consistent with the data shown in **Fig. 3a**. However, no difference in percentage or total cell numbers of ILC2s was noticed (**Supplementary Fig. 4f**). Nevertheless, GM-CSF produced by lung ILC2s was dramatically reduced due to Bhlhe40 deficiency, while there was no difference in the production of classical type 2 cytokines including IL-5, IL-13 and IL-4 (**Supplementary Fig. 4g-4h**). Among the mice chronically challenged with papain for 16 days, ILC2-specific *Bhlhe40* deficient mice did not show any reduction in eosinophils recruitment in lung compared to *Bhlhe40*^fl/fl^ mice (**Fig. 4c** and **Supplementary Fig. 4i**), which may be explained by a redundant effect of ILC2s and Th2 cells. Similarly, there was no difference in percentage and total cell numbers of ILC2s (**Supplementary Fig. 4j**). Even so, GM-CSF produced by lung ILC2s was reduced in the absence of Bhlhe40, while Bhlhe40 deletion in ILC2s did not alter the expression of classical type 2 cytokines (IL-5, IL-13 and IL-4) produced by ILC2s (**Supplementary Fig. 4k-4l**). Interestingly, IL-10 was increased in *Bhlhe40*^fl/fl^-*Klrg1*-Cre ILC2s compared with *Bhlhe40*^fl/fl^ ILC2s. Overall, Bhlhe40 was found to promote GM-CSF expression while inhibiting IL-10 expression. Bhlhe40 has been reported to repress IL-10 expression in Th1 and Th2 cells ^21, 22, 31^. Our findings here extend the important role of Bhlhe40 in regulating IL-10 from adaptive immune response to innate immune response.

**Fig. 4.**
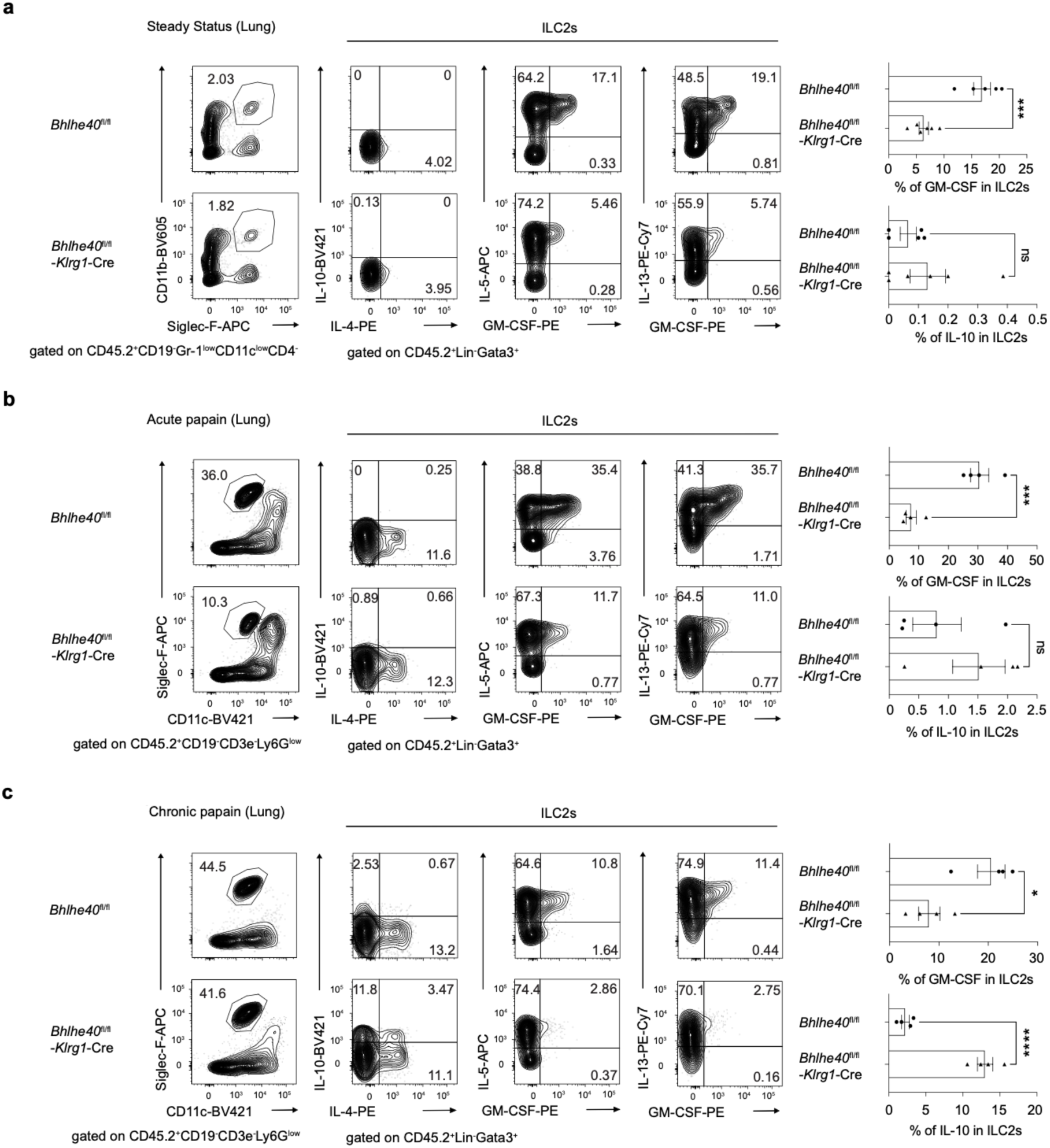
Bhlhe40 is required for GM-CSF but represses IL-10 in ILC2s. **a**, *Bhlhe40*^fl/fl^ and *Bhlhe40*^fl/fl^-*Klrg1*-Cre mice at steady state were analyzed. Left, eosinophils and cytokines production in lung ILC2s were shown by flow cytometry. Right, summary of percentage of GM-CSF and IL-10 in lung ILC2s. **b**, *Bhlhe40*^fl/fl^ and *Bhlhe40*^fl/fl^-*Klrg1*-Cre mice were acutely challenged with papain for 3 days and analyzed. Left, eosinophils and cytokines production in lung ILC2s were shown by flow cytometry. Right, summary of percentage of GM-CSF and IL-10 in lung ILC2s. **c**, *Bhlhe40*^fl/fl^ and *Bhlhe40*^fl/fl^-*Klrg1*-Cre mice were chronically challenged with papain for 16 days and analyzed. Left, eosinophils and cytokines production in lung ILC2s were shown by flow cytometry. Right, summary of percentage of GM-CSF and IL-10 in lung ILC2s. **a** show data from two independent experiments, each with at least one mouse for every genotype, with *Bhlhe40*^fl/fl^ (*n* = 5) and *Bhlhe40*^fl/fl^-*Klrg1*-Cre (*n* = 6) mice. **b** and **c**, show summarized results from three independent experiments with *Bhlhe40*^fl/fl^ (*n* = 4) and *Bhlhe40*^fl/fl^-*Klrg1*-Cre (*n* = 4) mice. * p < 0.05, ** p < 0.01, *** p < 0.001, **** p < 0.0001, Student’s *t*-test. Error bars indicate SEM.

### Differential regulation of cytokine production by Bhlhe40 in ILC2s and Th2 cells

Bhlhe40 has been reported to regulate the expression of type 2 cytokines in Th2 cells ^31^, however, we did not observe any role of Bhlhe40 in regulating these cytokines in ILC2s. To directly compare the role of Bhlhe40 in cytokine regulation in ILC2s and Th2 cells, we chronically challenged *Bhlhe40*^fl/fl^, *Bhlhe40*^fl/fl^-*Klrg1*-Cre, and *Bhlhe40*^fl/fl^-*Cd4*-Cre mice with papain for 16 days within the same experiment. Compared with *Bhlhe40*^fl/fl^ mice, there was no significant differences in eosinophil recruitment in mice with *Bhlhe40* deletion either in ILC2s (*Bhlhe40*^fl/fl^-*Klrg1*-Cre) or in T cells (*Bhlhe40*^fl/fl^-*Cd4*-Cre, **Fig. 5a** and **Fig. 5d**). And *Bhlhe40* deletion had no effect on the expansion and total cell numbers of ILC2s or Th2 cells (**Fig. 5b** and **Fig. 5c**). Nevertheless, Bhlhe40 deficiency increased IL-10 in both ILC2s and Th2 cells. However, while Bhlhe40 deficiency in ILC2s reduced their GM-CSF production, Bhlhe40 deficiency in Th2 cells decreased the frequency of the IL-13^+^IL-5^+^ population within Th2 cells (**Fig. 5b** and **Fig. 5c**), highlighting the distinct cytokine programs regulated by Bhlhe40 in these two cell types. To further analyze a possible redundant function of ILC2s and Th2 cells in eosinophil recruitment, we crossed *Bhlhe40*^fl/fl^-*Klrg1*-Cre mice with *Bhlhe40*^fl/fl^-*Cd4*-Cre mice to generate *Bhlhe40* double conditional knockout mice (*Bhlhe40*^fl/fl^-*Klrg1*-Cre-*Cd4*-Cre with Bhlhe40 deficiency in both ILC2s and Th2 cells). Indeed, 16 days after chronically papain challenging, compared with *Bhlhe40*^fl/fl^, eosinophils were dramatically reduced in lung from the mice with Bhlhe40 deficiency in both ILC2s and Th2 cells (**Fig. 5e** and **Fig. 5f**), indicating that there is a functional redundancy of ILC2s and Th2 cells on eosinophil recruitment regulated by Bhlhe40. Flow cytometry results shown in **Supplementary Fig. 5a** confirmed cytokine regulation by Bhlhe40 in ILC2s and Th2 cells from the *Bhlhe40*^fl/fl^-*Klrg1*-Cre-*Cd4*-Cre mice.

**Fig. 5.**
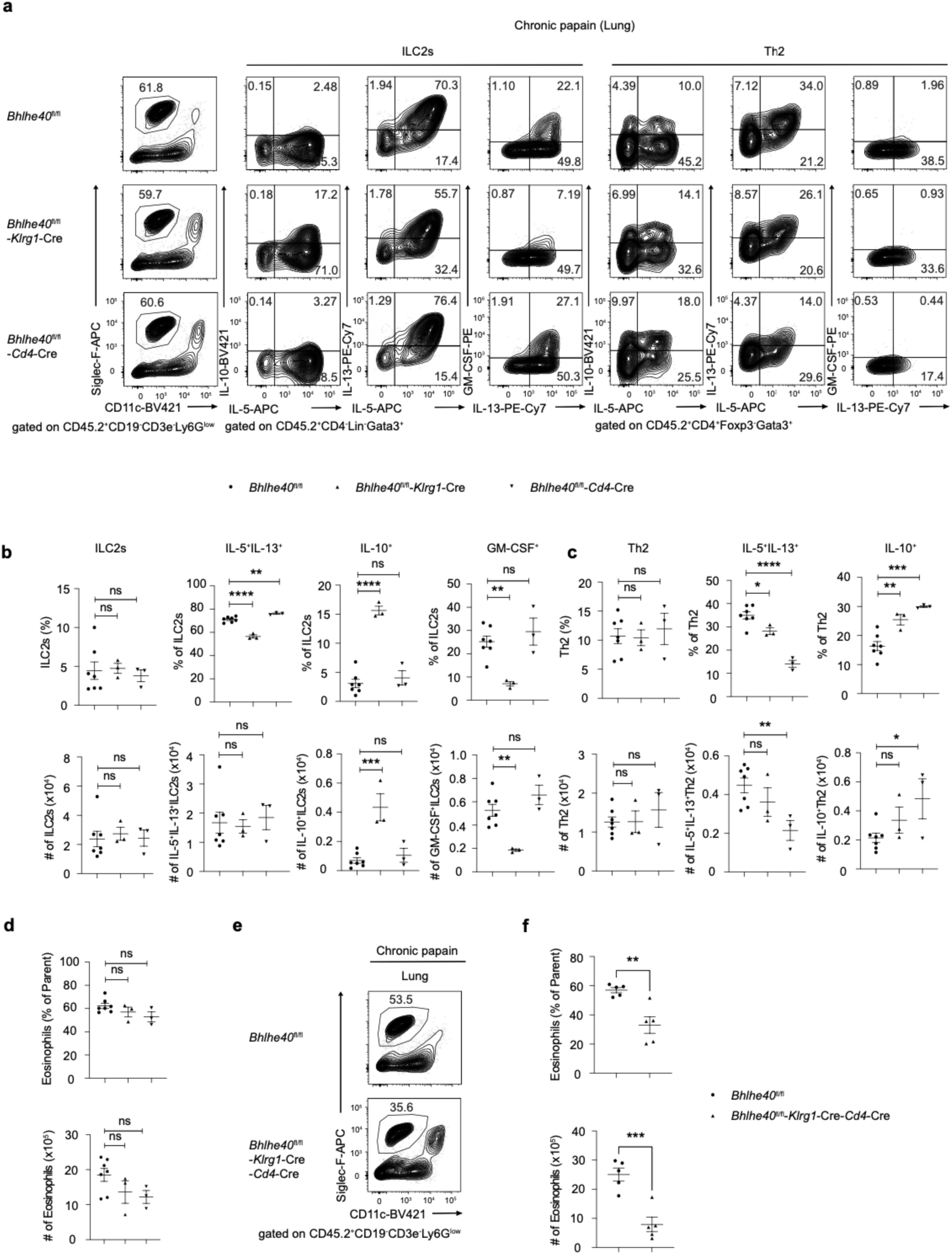
Bhlhe40 shows different functions in ILC2s and Th2 cells. **a**, *Bhlhe40*^fl/fl^, *Bhlhe40*^fl/fl^-*Klrg1*-Cre and *Bhlhe40*^fl/fl^-*Cd4*-Cre mice were chronically challenged with papain for 16 days, and lung were collected and analyzed for eosinophils recruitment, cytokines (IL-10, IL-5, IL-13 and GM-CSF) production in ILC2s and Th2 by flow cytometry. **b**, Percentage and cell number of ILC2s, cytokine-producing ILC2s (IL-10^+^, IL-5^+^IL-13^+^ and GM-CSF^+^) in lung were summarized from **a**. **c**, Percentage and cell number of Th2, cytokine-producing Th2 (IL-10^+^, IL-5^+^IL-13^+^) in lung were summarized from **a**. **d**, Percentage and cell number of eosinophils in lung were summarized from **a**. **e**, *Bhlhe40*^fl/fl^ and *Bhlhe40*^fl/fl^-*Klrg1*-Cre-*Cd4*-Cre mice were chronically challenged with papain for 16 days, and lung were collected and analyzed for eosinophils recruitment by flow cytometry. **f**, Percentage and cell number of eosinophils in lung were summarized from **e**. **a**-**d** show the combined results from two independent experiments with *Bhlhe40*^fl/fl^ (*n* = 7), *Bhlhe40*^fl/fl^-*Klrg1*-Cre (*n* = 3) and *Bhlhe40*^fl/fl^-*Cd4*-Cre (*n* = 3) mice. **e**-**f** show summarized results from two independent experiments with a total of *Bhlhe40*^fl/fl^ (*n* = 5), and *Bhlhe40*^fl/fl^-*Klrg1*-Cre-*Cd4*-Cre (*n* = 5) mice. * p < 0.05, ** p < 0.01, *** p < 0.001, **** p < 0.0001, Student’s *t*-test. Error bars indicate SEM.

### Single cell analyses of Bhlhe40-mediated gene regulation in ILC2s

Notably, we found that GM-CSF and IL-10 were mutual exclusively expressed by ILC2s from lung, BALF and mediastinal draining lymph nodes (dLNs, **Supplementary Fig. 5b**), suggesting that GM-CSF-and IL-10-producing ILC2s may represent distinct subsets of ILC2s. To get a broader picture on how Bhlhe40 regulates ILC2 functions, we isolated lung ILC2s from chronically papain challenged *Bhlhe40*^fl/fl^ mice (WT) and *Bhlhe40*^fl/fl^-*Klrg1*-Cre mice (KO), and performed single cell RNA-Seq and single cell ATAC-Seq (assay for transposase-accessible chromatin with sequencing) by using the 10x Genomics “multiomics” technology (**Fig. 6a**). The sorting strategy was shown in **Supplementary Fig. 6a**, and *Klrg1*-Cre was confirmed by GFP expression linked to iCre as expected based on the original designed ^32^. Five major clusters (0, 1, 2, 3 and 4) of ILC2s were identified from the scRNA-Seq data (**Fig. 6b**). Cluster specific signature genes were shown in **Fig. 6c** to separate different ILC2 clusters. Compared with WT, cluster 0 and cluster 2 ILC2 subpopulations were increased after *Bhlhe40* deletion, while clusters 1, 3, 4 subpopulations were decreased in *Bhlhe40* conditional knockout ILC2s (**Fig. 6d** and **Fig. 6e**). Our previous data showed that IL-10 was increased and GM-CSF was decreased at the protein level after *Bhlhe40* deletion in ILC2s. To confirm whether this regulation is at the mRNA level, we checked *Il10* and *Csf2* (gene encoding GM-CSF) expression in ILC2s using our scRNA-Seq datasets, and found the differences were consistent with their changes at the protein level (**Fig. 6f** and **Fig. 6g**). Since we performed scRNA-Seq and scATAC-Seq at the same time, we analyzed ILC2 clusters by integrating these two datasets and again showed five clusters of ILC2s (**Supplementary Fig. 6b**). Consistent with our previous findings, integrated analysis of the single-cell multimodal datasets showed that *Il10* expression was increased in *Bhlhe40*-deficient ILC2s whereas *Csf2* expression was reduced (**Supplementary Fig. 6c-6d**). Collectively, these single-cell multiomic data confirmed that Bhlhe40 positively regulates *Csf2* expression while repressing *Il10* expression at the transcriptional level in ILC2s. However, both IL-10 and GM-CSF can be produced by different clusters of ILC2s.

**Fig. 6.**
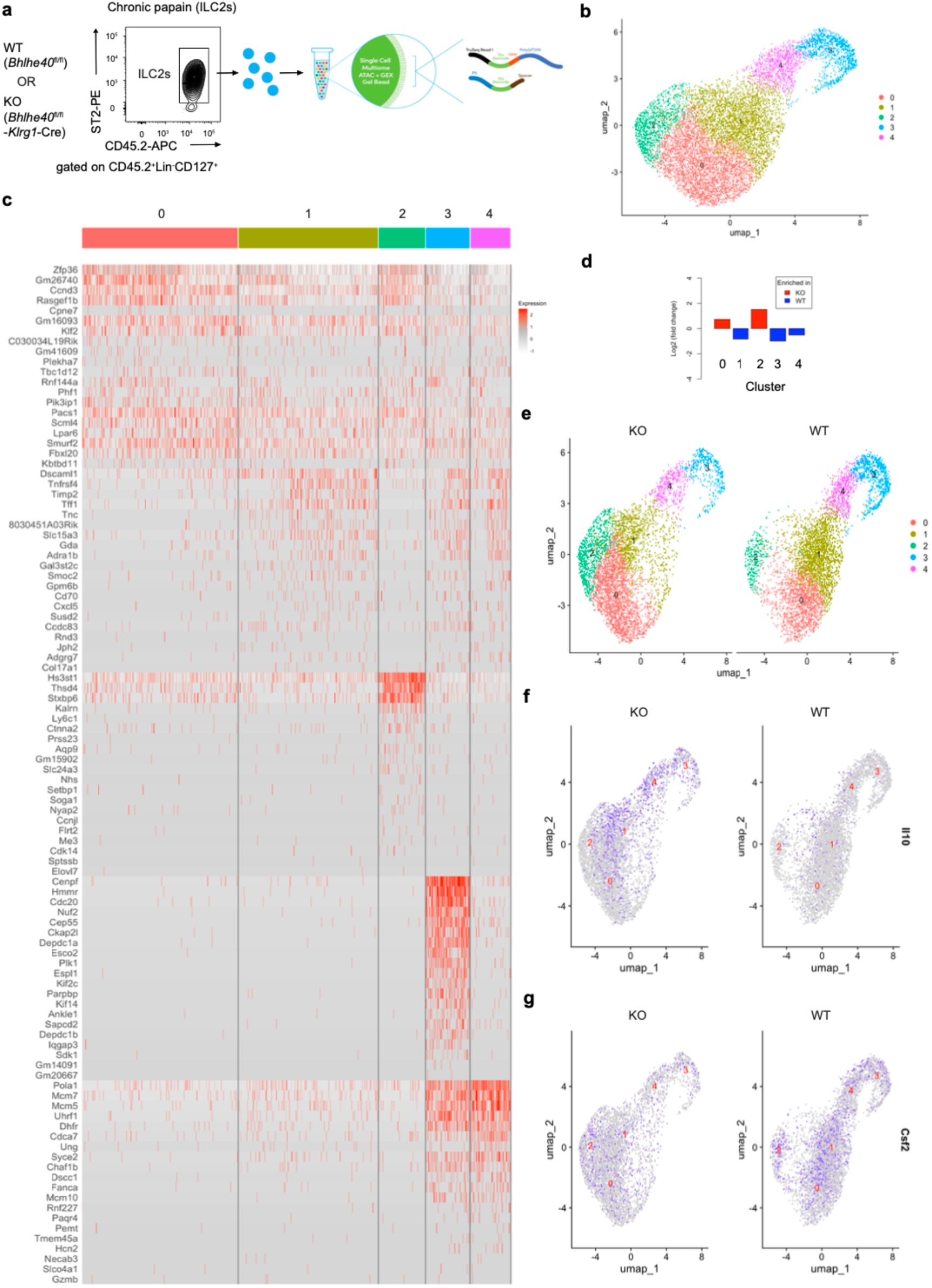
Multimodal datasets analysis of Bhlhe40’s function in ILC2s. **a**, WT (*Bhlhe40*^fl/fl^) and KO (*Bhlhe40*^fl/fl^-*Klrg1*-Cre) mice were chronically challenged with papain for 16 days. Cell suspension from lung was collected, stained and sorted for ILC2s (CD45.2^+^Lin^-^CD127^+^T1/ST2^+^), which were then processed for single-cell RNA-Seq and single-cell ATAC-Seq. **b**, five clusters of ILC2s were generated based on the scRNA-Seq results. **c**, Cluster specific signature genes were analyzed to distinguish 5 clusters (0, 1, 2, 3, 4). **d**, The difference in ILC2 clusters between WT and KO was plotted. **e**, Detailed differences of ILC2 clusters between WT and KO. **f**, Distribution of *Il10* expression in ILC2s clusters was compared between WT and KO. **g**, Distribution of *Csf2* expression in ILC2 clusters was compared between WT and KO. **a**-**g** show representative data from two independent experiments.

### Bhlhe40 directly regulates *Il10* and *Csf2* gene expression

To assess whether Bhlhe40 directly regulates *Il10* and *Csf2*, we performed anti-V5 ChIP-Seq with ILC2s from the Bhlhe40-V5 mice. WT and Bhlhe40-V5 mice were chronically challenged with papain for 16 days. Sorted ILC2s from lung were further cultured *in vitro* with cytokines (IL-2, IL-7 and IL-33) for one week to obtain enough cells to perform ChIP assay. ChIP-Seq data showed that Bhlhe40 directly bound to two elements (site 1 and site 2) at the *Il10* locus in ILC2s (**Fig. 7a**). These were the same regions that we have identified as Bhlhe40 binding elements in Th1 cells ^24^. Interestingly, chromatin accessibility at these two Bhlhe40-bound regions was increased following Bhlhe40 deletion in ILC2s as shown by our scATAC-Seq data (**Fig. 7a**, bottom), consistent with the idea that Bhlhe40 directly regulates *Il10* expression and gene accessibility by binding to its cis-regulatory elements. As shown in **Fig. 7b**, we also found that Bhlhe40 bound to the promoter of the *Csf2* gene in ILC2s, however, there was no changes in gene accessibility at this site after Bhlhe40 deletion (**Fig. 7b**), indicating that Bhlhe40 may regulate *Csf2* expression without affecting its gene accessibility.

**Fig. 7.**
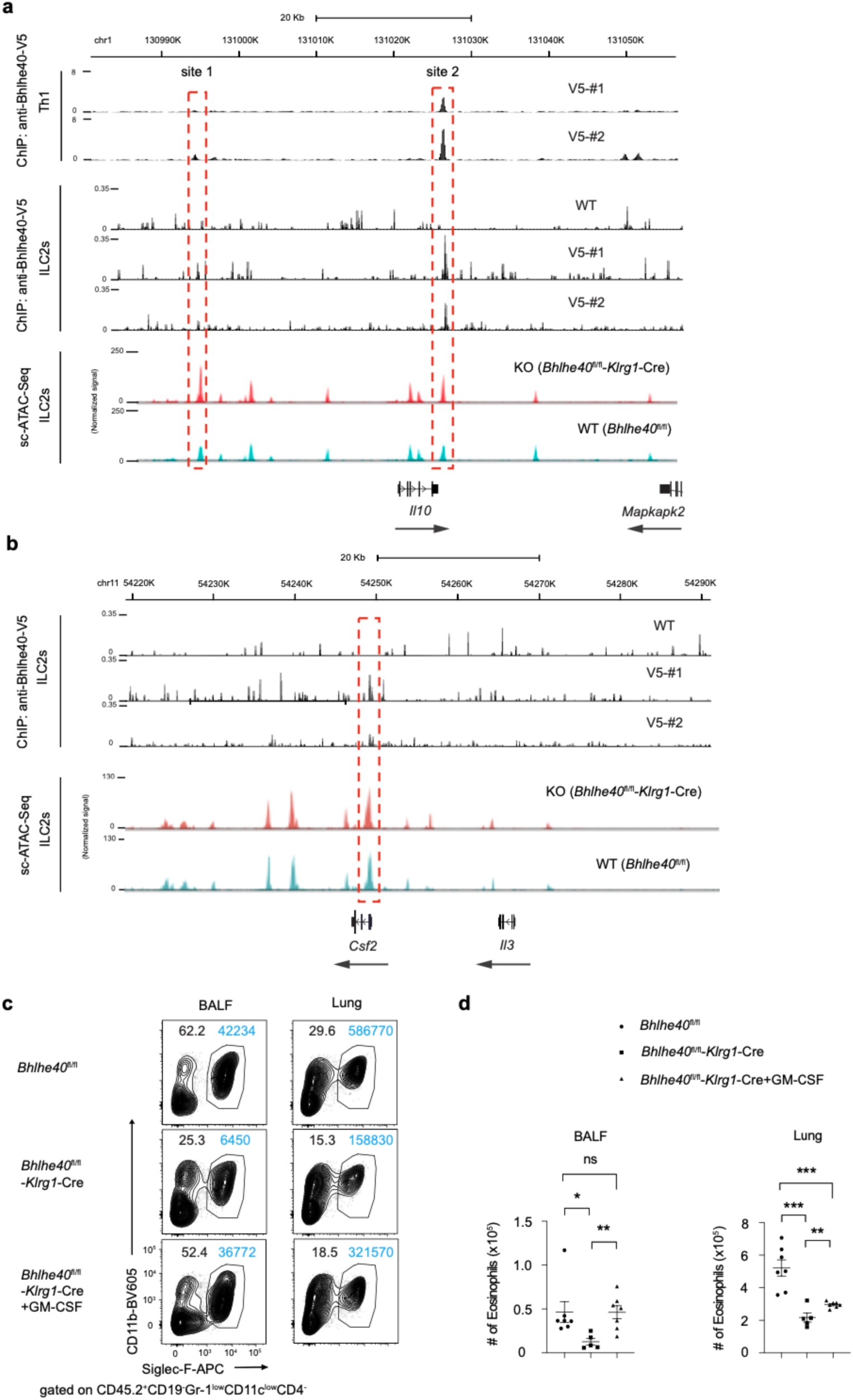
Bhlhe40 regulates ILC2 functions by directly repressing Il10 and inducing Csf2. **a**-**b**, WT and Bhlhe40-V5 mice were chronically challenged with papain for 16 days, and lung were harvested for staining and sorting ILC2s, which were later expanded in vitro for one week to get enough cell number for performing ChIP-Seq with anti-V5. **a**, *Il10* locus was shown in analyzing anti-Bhlhe40-V5 ChIP-Seq in ILC2s and Single-cell ATAC-Seq in ILC2s shown in Fig. 6. Anti-Bhlhe40-V5 ChIP-Seq data in Th1 were re-analyzed as a comparison. **b**, *Csf2* locus was shown in analyzing anti-Bhlhe40-V5 ChIP-Seq in ILC2s and Single-cell ATAC-Seq in ILC2s shown in Fig. 6. **c**, *Bhlhe40*^fl/fl^ and *Bhlhe40*^fl/fl^-*Klrg1*-Cre mice were pre-treated with PBS or GM-CSF for 3 days, and then acutely challenged with papain, together with PBS or GM-CSF for another 3 days. BALF and lung were collected and analyzed for eosinophils (CD45.2^+^CD19^-^ Gr-1^low^CD11c^low^CD4^-^Siglec-F^+^CD11b^+^) recruitment by flow cytometry. **d**, Cell number of eosinophils in BALF and lung were summarized from **c**. **a**-**b** show ChIP-Seq data from one experiment with two biological replicates, and sc-ATAC-Seq representative data from two independent experiments. **c**-**d** show summarized results from two independent experiments with *Bhlhe40*^fl/fl^ (*n* = 7), *Bhlhe40*^fl/fl^-*Klrg1*-Cre (*n* = 5) and *Bhlhe40*^fl/fl^-*Klrg1*-Cre + GM-CSF (*n* = 7) mice. * p < 0.05, ** p < 0.01, *** p < 0.001, **** p < 0.0001, Student’s *t*-test. Error bars indicate SEM.

Finally, to prove whether the reduction in GM-CSF production in ILC2s caused by Bhlhe40 deficiency is responsible for reduced eosinophil recruitment, we performed a rescue experiment. As shown in **Fig. 7c** and **Fig. 7d**, administration of GM-CSF fully rescued the reduction of eosinophils in the BALF caused by Bhlhe40 deficiency in ILC2s, and it also partially rescued eosinophil recruitment in the lung. In summary, we propose a working model of Bhlhe40-mediated gene regulation in ILC2s and in Th2 cells shown in **Supplementary Fig. 7**. Collectively, our findings identify Bhlhe40 as a key transcriptional regulator of ILC2 effector functions by promoting GM-CSF expression while repressing IL-10. We further demonstrate that Bhlhe40-depedent GM-CSF production, together with IL-5, is critical for eosinophil recruitment during ILC2-driven type 2 immune responses.

## DISCUSSION

ILC2s share functional characteristics with Th2 cells but respond much more rapidly and play a critical role during type 2 innate immune responses ^33^. GATA3, the master transcription factor governing both Th2 cells and ILC2s, regulates the expression of a series of genes required for lineage commitment and functions in both cell types ^9^. However, how other transcription factors are involved in Th2 and ILC2 biology remains unclear. Bhlhe40 has been reported to regulate Th2 cell-mediated responses during type 2 immunity ^31^; nevertheless, the role of Bhlhe40 in ILC2s has not been investigated. Our study demonstrates that Bhlhe40 is required for ILC2 effector functions. Furthermore, we show that Bhlhe40 regulates distinct transcriptional programs in ILC2s and Th2 cells. Consistent with the previous report ^31^, Bhlhe40 promotes the expression of classical type 2 cytokines (IL-4, IL-5 and IL-13) in Th2 cells. In contrast, Bhlhe40 did not regulate these cytokines in ILC2s but instead promoted GM-CSF production, thereby contributing to eosinophil recruitment and type 2 lung inflammation. These findings reveal a previously unrecognized, cell type-specific function of Bhlhe40 in coordinating cytokine programs during type 2 immune responses.

IL-5 produced by ILC2s or Th2 cells is well-known as the key cytokine recruiting eosinophils to sites of inflammation during type 2 immune responses ^13^. However, while both ILC2s and Th2 cells can produce IL-5, ILC2s are more efficient in recruiting eosinophils than Th2 cells suggesting that there may be another mechanism in addition to IL-5 production contributing to eosinophil recruitment by ILC2s. By using multiple Bhlhe40 deficient mice, we found that eosinophil recruitment was severely reduced with Bhlhe40 deficiency in ILC2s, and this was associated with a reduction of GM-CSF but not IL-5 production by Bhlhe40 deficient ILC2s. Importantly, eosinophil recruitment in these mice can be rescued by administration of exogenous GM-CSF. Our findings highlight an efficient regulatory pathway for eosinophils recruitment involving GM-CSF production by ILC2s. Since GM-CSF is well known to function in multiple cell types and to regulate immune-cell development and activation ^14, 15^, it is of interest to determine whether ILC2-derived GM-CSF contributes to the generation or functional regulation of these cells.

IL-1β and TCR signaling have been reported to induce Bhlhe40 expression in T cells ^34, 35^. However, the signaling pathways responsible for Bhlhe40 induction during ILC2 activation remain unknown. The activation of ILC2s is regulated by epithelial-derived alarmins, including IL-33, IL-25, and TSLP, as well as lipid mediators and neuroimmune signals ^36, 37^. Since IL-1β can induce Bhlhe40 in T cells, it is likely that IL-33 (a cytokine of the IL-1 superfamily) may induce Bhlhe40 expression in ILC2s, which requires further investigation.

Bhlhe40 has been reported to promote inflammatory responses in Th1 cells by enhancing IFN-γ production while repressing IL-10 expression ^20, 21, 22^. In ILC2s, we similarly found that Bhlhe40 promotes pro-inflammatory functions by inducing GM-CSF production and suppressing IL-10 expression. Therefore, there is a conserved regulatory mechanism between innate and adaptive lymphocytes involving Bhlhe40 to control appropriate inflammation.

Although Bhlhe40 regulates classical type 2 cytokine production in Th2 cells, it does not have the same function in ILC2s, suggesting that distinct regulatory mechanisms may exist between these innate and adaptive type 2 lymphocytes. In addition, while ILC2s can produce GM-CSF, we did not observe GM-CSF production by Th2 cells in our in vivo models of type 2 immune responses. It is possible that Bhlhe40 may interact with different partners in different cell types. Proteomic approaches such as mass spectrometry may allow us to identify distinct Bhlhe40-interacting partners in Th2 cells and ILC2s in the future. GATA3 has been reported to promote GM-CSF production in Th17 ^38^, raising the possibility that Bhlhe40-dependent regulation of GM-CSF in ILC2s may also involve GATA3. However, since both ILC2 and Th2 cells express Bhlhe40 and GATA3, additional factors must be involved in the differential regulation of GM-CSF.

Bhlhe40 expression has been reported in CD4^+^ T cells from patients with rheumatoid arthritis ^39^. Given the increasing incidence of severe respiratory viral infections, particularly in immunodeficient patients with impaired viral clearance, it would be valuable to investigate whether dysregulated Bhlhe40 expression or mutation contributes to defective immune responses in these patients. In this context, therapeutic strategies involving GM-CSF may merit further consideration. Given that GM-CSF has been approved by the FDA for clinical applications such as hematopoietic stem cell transplantation and immune reconstitution ^14, 40^, a potential side effect of GM-CSF in causing eosinophilia should also be considered.

In conclusion, our study identifies Bhlhe40 as a key regulator of the pro-inflammatory functions of ILC2s. By repressing IL-10 while promoting GM-CSF production, Bhlhe40 enhances eosinophil recruitment to sites of inflammation and thereby contributes to type 2 immune responses. Although Bhlhe40 is expressed in both ILC2s and Th2 cells, it regulates distinct cytokine programs in these two cell types. In ILC2s, Bhlhe40 is required for GM-CSF production while repressing IL-10 expression, whereas in Th2 cells, Bhlhe40 promotes the production of the canonical type 2 cytokines IL-4, IL-5, and IL-13, while similarly repressing IL-10 expression. Thus, Bhlhe40 promotes eosinophil recruitment through distinct cell type-specific mechanisms: by inducing GM-CSF in ILC2s and by promoting type 2 cytokine production in Th2 cells. Together, these complementary functions contribute to the development of type 2 lung inflammation.

## ONLINE METHODS

### Mice

All the mouse strains are on the C57BL/6 background. Wide type (WT) C57BL/6 mice were purchased from Taconic Farms. *Bhlhe40*^fl/fl^ and *Bhlhe40*^fl/fl^-*Cd4*-Cre mice were previously generated in a previous report ^21^. Bhlh40-V5 tagged mice and *Bhlhe40* germline deficient mice were previously reported ^24^. *Bhlhe40*^fl/fl^-*Klrg1*-Cre mice were generated by crossing *Bhlhe40*^fl/fl^ mice with *Klrg1*-Cre mice ^32^, in which the Cre recombinase expression is under the control of *Klrg1* locus allowing ILC2-specific deletion of loxP-flanked *Bhlhe40* exon 4. *Bhlhe40*^fl/fl^-*Klrg1*-Cre-*Cd4*-Cre mice were generated by crossing *Bhlhe40*^fl/fl^-*Klrg1*-Cre mice with *Bhlhe40*^fl/fl^-*Cd4*-Cre mice. All experiments used mice at 6–16 weeks of age under an animal study protocol (ASP: LISB-8E) approved by the National Institute of Allergy and Infectious Diseases Animal Care and Use Committee.

### Papain-induced airway inflammation

During papain-induced airway inflammation, mice were anesthetized with isoflurane and administered with 40 μg of papain in 20 μl of phosphate-buffered saline (PBS) intranasally (*i.n.*) on days 0, 1, and 2. For acute papain studies, mice were euthanized on day 3. For chronic papain studies, mice were allowed to rest for 10 days and then rechallenged with papain on days 13, 14, and 15. Mice were euthanized 24 h after the final *i.n.* challenge. Bronchoalveolar lavage fluid (BALF) and lungs were collected for subsequent analyses.

### Pulmonary GM-CSF aspiration

Mice were anesthetized using isoflurane and 25 μL of GM-CSF (1 μg) dissolved in PBS was administered by pharyngeal aspiration as previously described ^41^. This treatment was administered once daily for three consecutive days. Mice were then challenged once daily for an additional three days with papain (40 μg) together with GM-CSF (1 μg). 24 h after the final challenge, mice were euthanized, and BALF and lungs were collected for downstream analyses.

### Nippostrongylus brasiliensis infection

Mice were subcutaneously (*s.c.*) inoculated with 500 infective third-stage larvae (L3) of *Nippostrongylus brasiliensis* (*N. brasiliensis*). The adult worms in the intestine was performed and counted as previously described ^42^. Lungs, BALF, and mesenteric lymph nodes (mLNs) were collected for further analyses.

### Cell isolation

BALF was obtained by flushing lung twice with 0.8 mL of PBS using a syringe cannula. The lung lobes were chopped into small pieces and digested with Liberase (20 μg/ml) and deoxyribonuclease I (DNase I) (10 U/ml) in RPMI-1640 supplemented with 3% fetal bovine serum (FBS) for 20 min at 37°C. The digestion was terminated by adding three times ice-cold RPMI 1640 + 3% FBS. Tissue debris were removed by filtration through a 40-μm cell strainer, and red blood cells were lysed using ACK lysis buffer. The draining (mediastinal) lymph nodes (dLNs) were mechanically dissociated and passed through 40-μm cell strainers to obtain single-cell suspensions.

Small intestines were collected from euthanized mice, emptied of luminal contents, cleared of external fat, and stripped of Peyer’s patches. The intestines were opened longitudinally and cut into 1-cm pieces. Tissue fragments were subjected to a two-step digestion protocol. First, samples were incubated for 20 min at 37°C in RPMI-1640 medium supplemented with 3% FBS, 1 mM EDTA, and 1 mM dithiothreitol (DTT). The tissues were then filtered through gauze, washed with PBS, minced, and further digested for 45 min at 37°C in RPMI-1640 medium containing 20 μg/mL Liberase and 10 U/mL DNase I. Following digestion, the cell suspension was passed through a 40-μm cell strainer. Residual tissue fragments were mechanically dissociated and washed through the strainer. Cells were collected by centrifugation at 1,600 rpm for 6 min, and lymphocytes were enriched using 40% Percoll. Purified lymphocytes were subsequently used for staining and flow cytometric analysis. Mesenteric lymph nodes (mLNs) were processed in the same as with dLNs.

### Flow cytometry and cell sorting

Single cell suspensions were either stained directly or stimulated with PMA (phorbol 12-myristate 13-acetate) and ionomycin in the presence of GolgiStop reagent (monensin) for 4 h at 37°C prior to staining. Briefly, cells were first incubated with anti-CD16/32 antibody (clone 2.4G2) at 4°C for 15 min to block Fc receptors. Surface markers were then stained with fluorochrome-conjugated antibodies in PBS containing 3% FBS for 30 min at room temperature. For intracellular staining of transcription factors, cells were fixed and permeabilized using the Foxp3 Staining Buffer Set (00-5523-00, eBioscience) according to the manufacturer’s instructions. For intracellular cytokine staining in combination with transcription factor staining, cells were first fixed with 1% paraformaldehyde (PFA) for 5 min at room temperature, washed, and then processed using the same fixation and permeabilization protocol. Flow cytometry data were acquired using LSRFortessa or FACSymphony flow cytometers (BD Biosciences) and analyzed with FlowJo software (Tree Star). Eosinophils were identified as CD45^+^CD3e^-^CD19^-^Ly6G^low^CD11c^low^Siglec-F^+^CD11b^+^; Th2 cells were identified as CD45^+^CD4^+^Foxp3^-^GATA3^+^; ILC2s were identified as CD45^+^CD4^-^ Lin^-^GATA3^+^. Cells were sorted on an FACS Aria cell sorter (BD Biosciences). To sort ILC2s from lung, cells were depleted with Lin antibodies (against CD3, CD5, CD19, γδTCR, CD11b, CD11c, NK1.1, FceR1, TER119, Ly6G/6C) before sorting and then identified as CD45^+^Lin^-^ CD127^+^T1/ST2^+^.

### scRNA-Seq and scATAC-Seq analysis

ILC2s were sorted from lung of mice chronically challenged with papain and prepared for Chromium Next GEM Single Cell Multiome ATAC + Gene Expression following the 10× protocol for nuclei isolation with a lysis time of three minutes. Nuclei were loaded onto the 10x Chromium controller to target 3 × 10^3^ to 5 × 10^3^ nuclei. Ctrl (*Bhlhe40*^fl/fl^) and KO (*Bhlhe40*^fl/fl^-*Klrg1*-Cre) ILC2s were captured separately in the same experiment, and two biological replicates of each genotype were captured in two independent experiments. Libraries were constructed according to the manufacturer’s instructions and sequenced on Illumina NextSeq 2000. For gene expression, the run parameters were 28×10×10×90, and for ATAC 50×8×24×49. The resulting sequencing data were processed using the 10x Genomics Cell Ranger ARC pipeline. Briefly, raw base call (BCL) files were demultiplexed into FASTQ files, aligned to the mouse reference genome (mm10), and processed to identify valid cell barcodes and unique molecular identifiers (UMIs). Gene expression and chromatin accessibility matrices were generated for downstream analyses, including quality control, normalization, dimensionality reduction, cell clustering, differential gene expression, and chromatin accessibility analyses using the Seurat ^43^ and Signac ^44^ packages in R.

### ChIP-Seq analysis

For ChIP-Seq analysis of Bhlhe40 binding in ILC2s, lung ILC2s were sorted from WT or Bhlhe40-V5 mice chronically challenged papain. Chromatin immunoprecipitation was performed using an anti-V5 antibody (R96025, Thermo Fisher Scientific). Cells were cross-linked with 1% formaldehyde for 10 min at room temperature and sonicated in shearing buffer (0.4% SDS in TE buffer) to generate DNA fragments ranging from 100 to 500 bp. Ten percent of the total chromatin was reserved as input control, and the remaining chromatin was subjected to immunoprecipitation. DNA–protein complexes were immunoprecipitated overnight in RIPA buffer with the anti-V5 antibody. DiaMag anti-mouse IgG-coated magnetic beads or DiaMag Protein A-coated magnetic beads were then added and incubated for an additional 4 h. Following extensive washing, cross-links were reversed, and DNA was purified. Purified DNA fragments (100–500 bp) were used to prepare indexed sequencing libraries, which were subsequently subjected to high-throughput sequencing. ChIP-seq data were processed using a standard analysis pipeline. Briefly, raw sequencing reads underwent base calling, demultiplexing, and quality assessment, followed by adapter trimming and quality filtering. Filtered reads were aligned to the mouse reference genome (mm10) using Bowtie2 ^45^, and alignment files were processed using SAMtools ^46^. Post-alignment quality metrics were evaluated before peak calling, visualization of genome-wide binding profiles, and downstream comparative analyses. ChIP-seq peaks were identified using MACS3 ^47^ with the appropriate control samples. Peak annotation, de novo motif enrichment, differential binding analysis, and overlap analysis were performed using HOMER ^48^. BAM files were converted to BED format using BEDTools ^49^, and normalized bigWig files were generated for visualization in the UCSC Genome Browser ^50^. Genome browser tracks were used to examine Bhlhe40 occupancy at selected genomic loci.

### Statistics

Samples were compared with Prism 10 software (GraphPad) by a two-tailed unpaired Student’s t-test. Data were presented as mean ± SEM. A p-value <0.05 was considered statistically significant and indicated as *; p < 0.01 was indicated as **; p < 0.001 was indicated as ***; and p < 0.0001 was indicated as ****. Not statistically significant was indicated as ns.

### Accession numbers and materials availability

The ChIP-Seq and single-cell multiomic (scRNA-Seq and scATAC-Seq) datasets have been deposited and are available at the Gene Expression Omnibus (GEO) database under the accession numbers: GSE342543, GSE342380 and GSE342381. Mouse strains generated in this study are available upon request to JZ after completion of a materials transfer agreement (MTA) with NIAID.

## ACKNOWLEDGEMENTS

We thank Aida Romero-Garcia and the NIAID Flow Cytometry Section for cell sorting, NIAID animal facilities for helping maintain our mouse strains, and Yan Xin and Yicheng Wang for pharyngeal aspiration technique help. Sequencing was performed with the Frederick National Lab for Cancer Research CCR Sequencing Facility. This research was supported by the Intramural Research Program of the National Institutes of Health (NIH). The contributions of the NIH author(s) made as part of their official duties as NIH federal employees are in compliance with agency policy requirements and are considered works of the US Government. However, the findings and conclusions presented in this paper are those of the authors and do not necessarily reflect the views of the NIH or the US Department of Health and Human Services.

## AUTHOR CONTRIBUTIONS

Conceptualization: XZ, JZ

Methodology: XZ, JS, YC, JN, XC

Investigation: XZ, DW, RKG, DP

Funding acquisition: JZ, RB, KZ

Supervision: JZ, RB, KZ

Writing – original draft: XZ, JZ

Writing – review & editing: XZ, JS, JN, RB, KZ, JZ

---

**SM Fig. 1.**
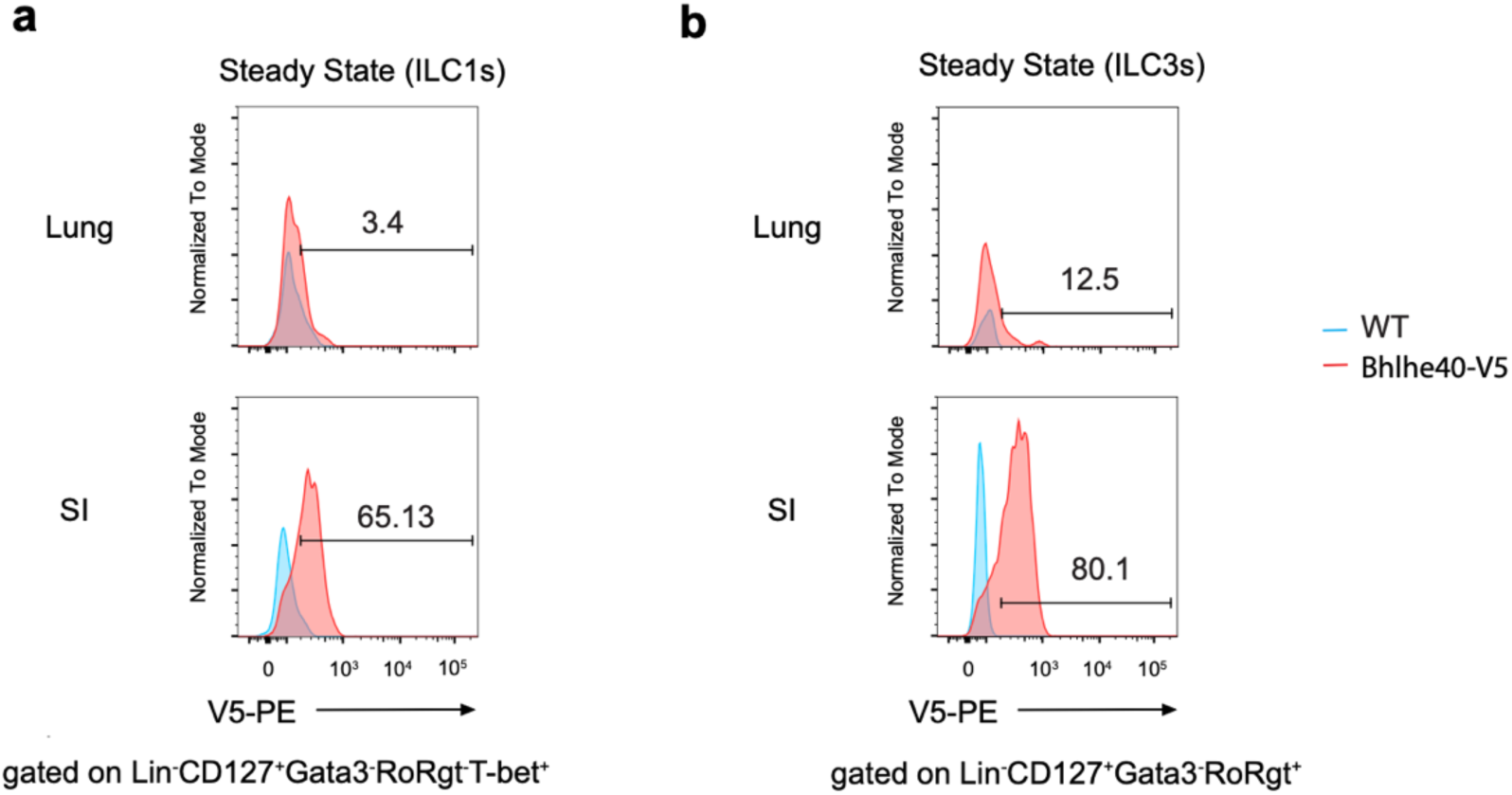
(related to Fig. 1) Bhlhe40 is expressed by gut ILC1s and ILC3s a, In steady state, Bhlhe40 expression in ILC1s (Lin^-^CD127^+^GATA3^-^RORγt^-^T-bet^+^) was indicated by anti-V5 staining in lung and small intestine (SI). **b**, In steady state, Bhlhe40 expression in ILC3s (Lin^-^CD127^+^GATA3^-^RORγt^+^) was indicated by anti-V5 staining in lung and small intestine. **a**-**b** show representative data from two independent experiments.

**SM Fig. 2.**
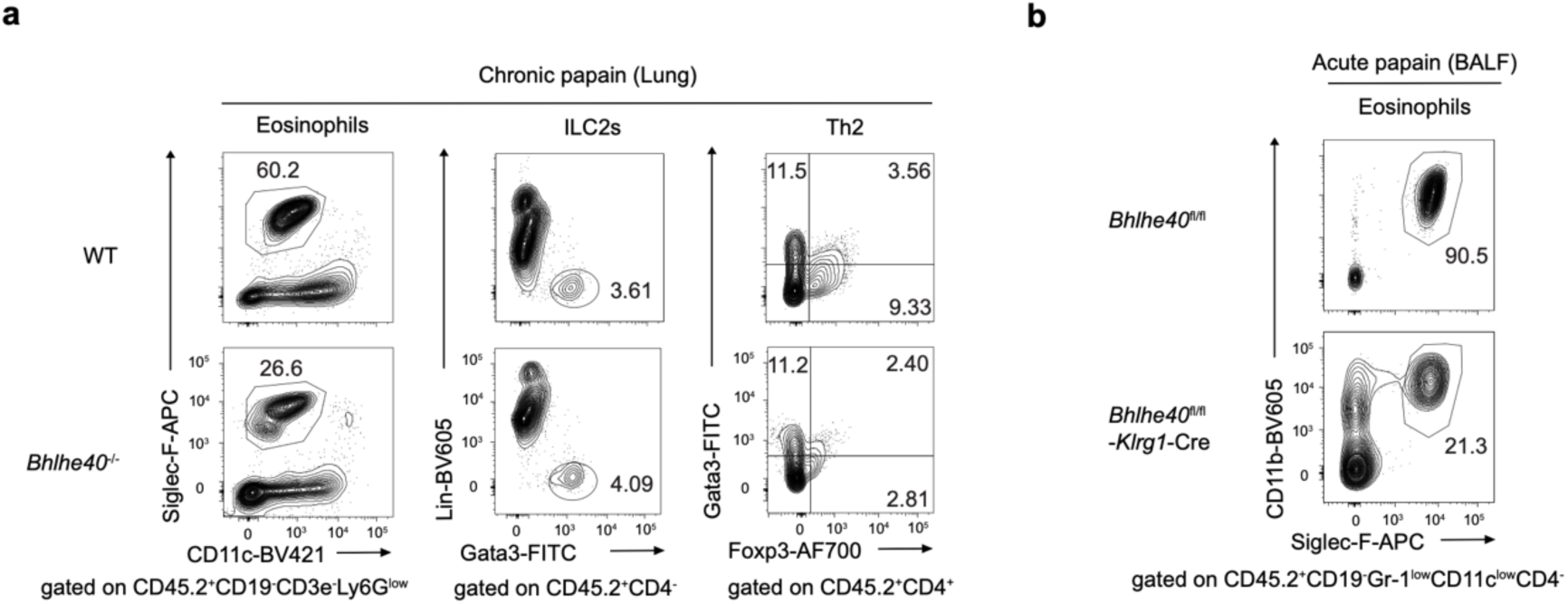
(related to Fig. 2 and Fig. 3) Germline or ILC2-specific Bhlhe40 deficient mice have reduced recruitment of eosinophils during type 2 immune responses a, WT and *Bhlhe40*^-/-^ mice were chronically challenged with papain for 16 days. Eosinophils (CD45.2^+^CD19^-^CD3e^-^Ly6G^low^CD11c^low^Siglec-F^+^), ILC2s (CD45.2^+^CD4^-^Gata3^+^Lin^-^) and Th2 (CD45.2^+^CD4^+^GTAT3^+^ Foxp3^-^) in lung were analyzed by flow cytometry. **b**, *Bhlhe40*^fl/fl^ and *Bhlhe40*^fl/fl^-*Klrg1*-Cre mice were acutely challenged with papain for 3 days. Eosinophils (CD45.2^+^CD19^-^Gr-1^low^CD11c^low^CD4^-^Siglec-F^+^CD11b^+^) in BALF were analyzed by flow cytometry. **a**-**b** show representative data from two independent experiments.

**SM Fig. 3.**
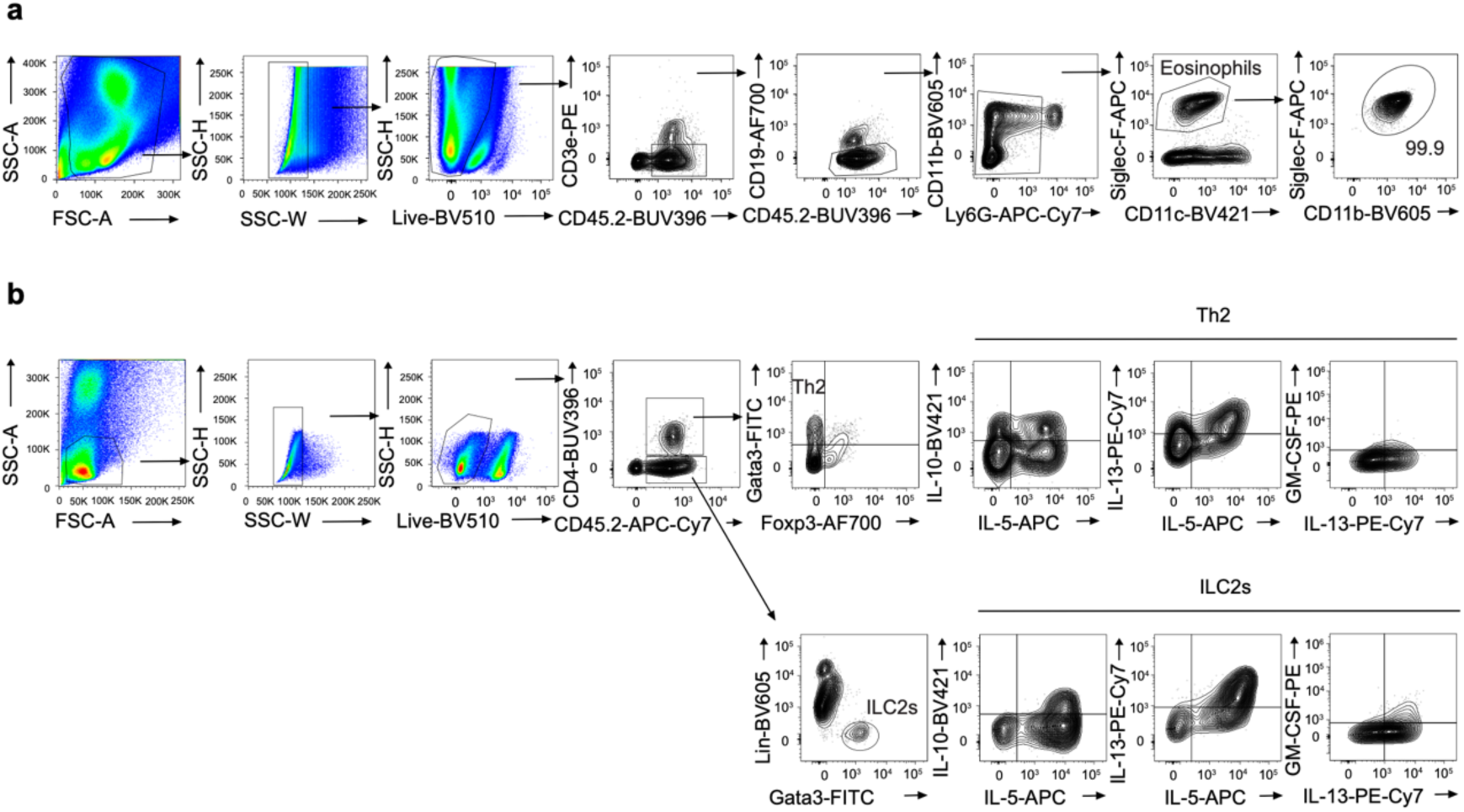
(related to Fig. 4 and Fig. 5)Gating strategy for flow cytometric analysis of eosinophils, ILC2s, Th2, and their cytokine production. WT or *Bhlhe40*^fl/fl^-*Klrg1*-Cre mice were chronically challenged with papain for 16 days. The lung cell suspension was stained and analyzed by flow cytometry. **a**, Gating strategy for identifying eosinophils. **b**, Gating strategy for identifying ILC2s, Th2, and cytokines (IL-10, IL-5, IL-13, and GM-CSF) produced in ILC2s or Th2.

**SM Fig. 4.**
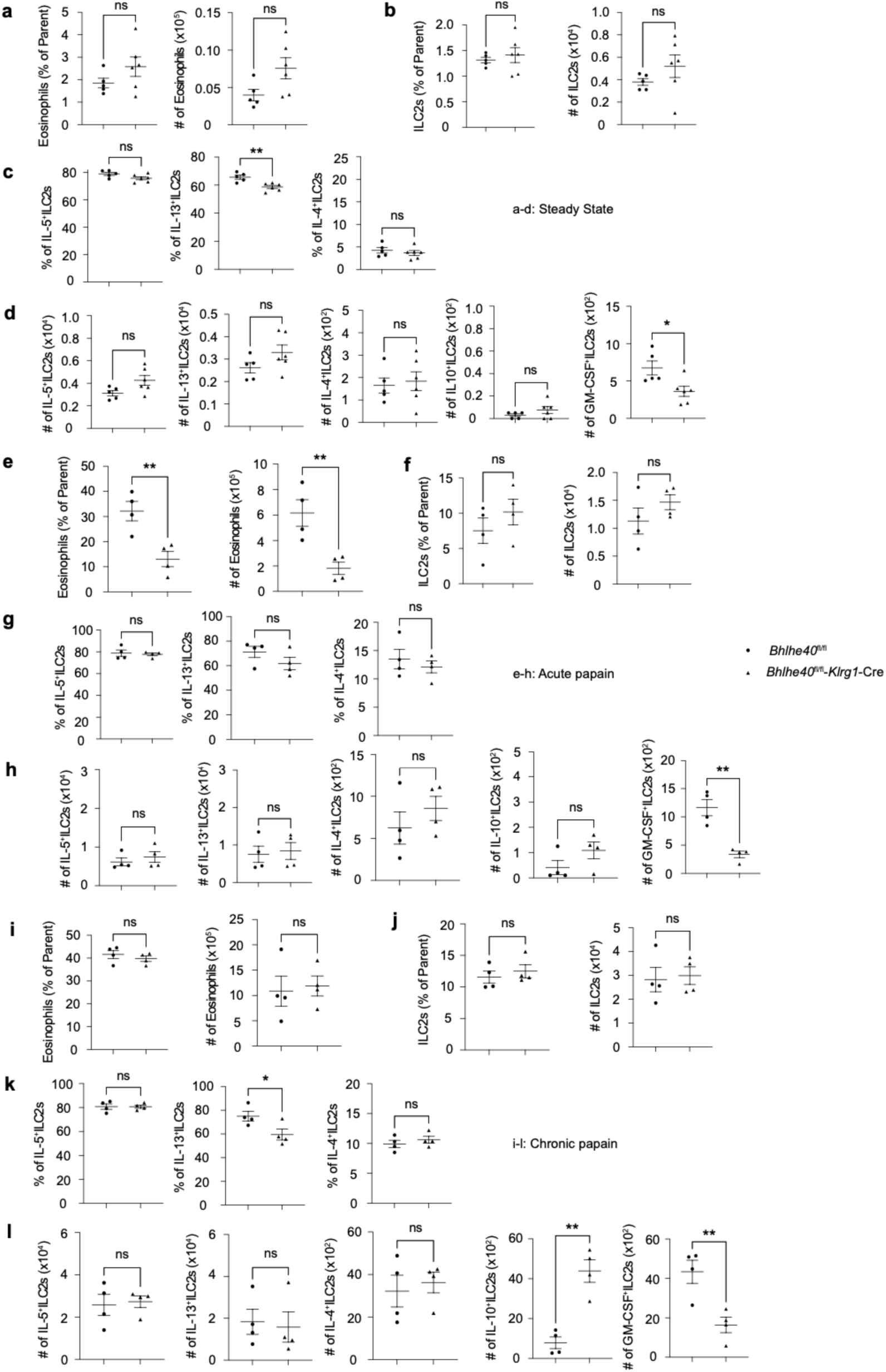
(related to Fig. 4) Bhlhe40 promotes GM-CSF but represses Il10 expression in ILC2s. Summary of percentage and cell number of eosinophils, ILC2s and cytokines produced by ILC2s in steady state (**a-d**), in acutely challenged with papain (**e-h**), and in chronically challenged with papain (**i-l**). **a**-**d** show summarized results from two independent experiments with *Bhlhe40*^fl/fl^ (*n* = 5) and *Bhlhe40*^fl/fl^-*Klrg1*-Cre (*n* = 6) mice. **e**-**l** show summarized results from three independent experiments with *Bhlhe40*^fl/fl^ (*n* = 4) and *Bhlhe40*^fl/fl^-*Klrg1*-Cre (*n* = 4) mice. * p < 0.05, ** p < 0.01, *** p < 0.001, **** p < 0.0001, Student’s *t*-test. Error bars indicate SEM.

**SM Fig. 5.**
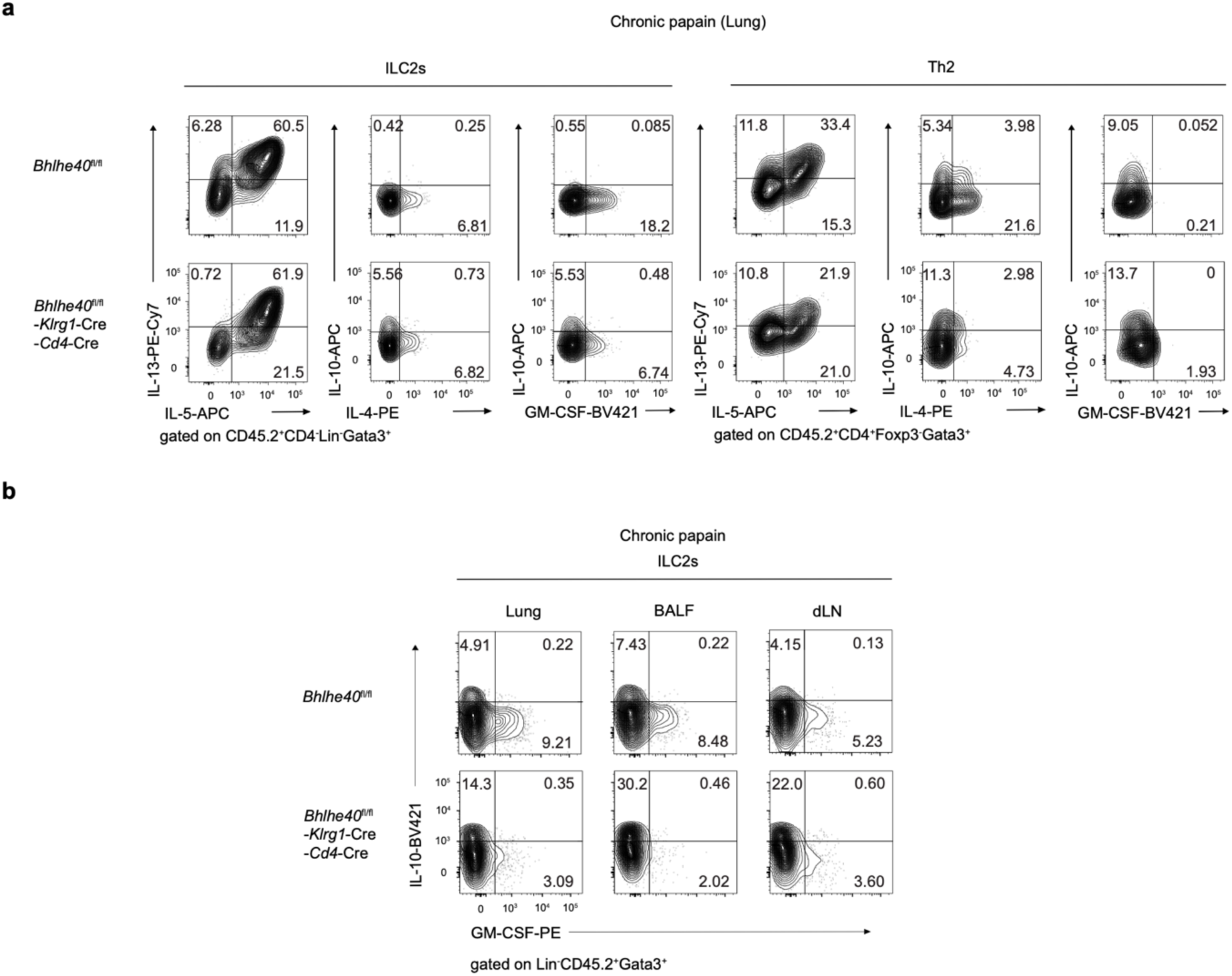
(related to Fig. 5) Bhlhe40 exhibits different functions in ILC2s and Th2 cells a, *Bhlhe40*^fl/fl^ and *Bhlhe40*^fl/fl^-*Klrg1*-Cre-*Cd4*-Cre mice were chronically challenged with papain for 16 days, and cell suspension from lung was stained and analyzed for cytokines (IL-5, IL-13, IL-4, IL-10, and GM-CSF) produced by ILC2s or Th2 cells by flow cytometry. **b**, *Bhlhe40*^fl/fl^ and *Bhlhe40*^fl/fl^-*Klrg1*-Cre mice were chronically challenged with papain for 16 days, and cell suspension from lung, BALF and dLN (draining-mediastinal lymph nodes) were stained and analyzed for cytokines (IL-10, and GM-CSF) produced by ILC2s by flow cytometry. **a**-**b** show representative data from two independent experiments.

**SM Fig. 6.**
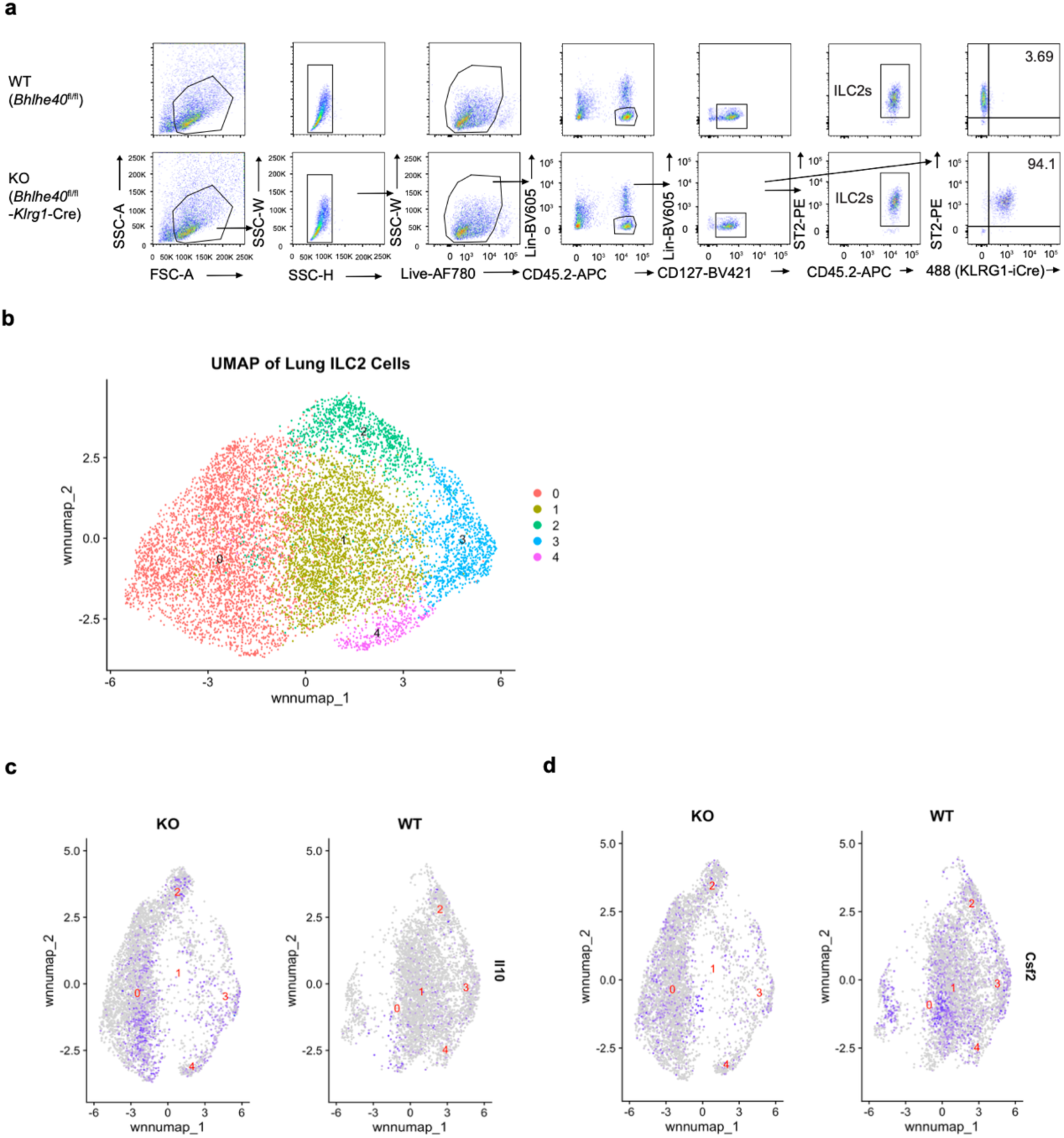
(related to Fig. 6) Multimodal datasets analyses of Bhlhe40’s functions in ILC2s a, Gating strategy for sorting ILC2s in lung cell suspension after chronically challenging WT (*Bhlhe40*^fl/fl^) and KO (*Bhlhe40*^fl/fl^-*Klrg1*-Cre) mice with papain for 16 days. **b,** Integrated single-cell RNA-seq data and single-cell ATAC-seq data to generate clusters of lung ILC2s. **c,** *Il10* distribution in ILC2 clusters was compared between WT and KO from the integrated data analysis. **d,** *Csf2* distribution in ILC2 clusters was compared between WT and KO from the integrated data analysis. **a**-**d** show representative data from two independent experiments.

**SM Fig. 7.**
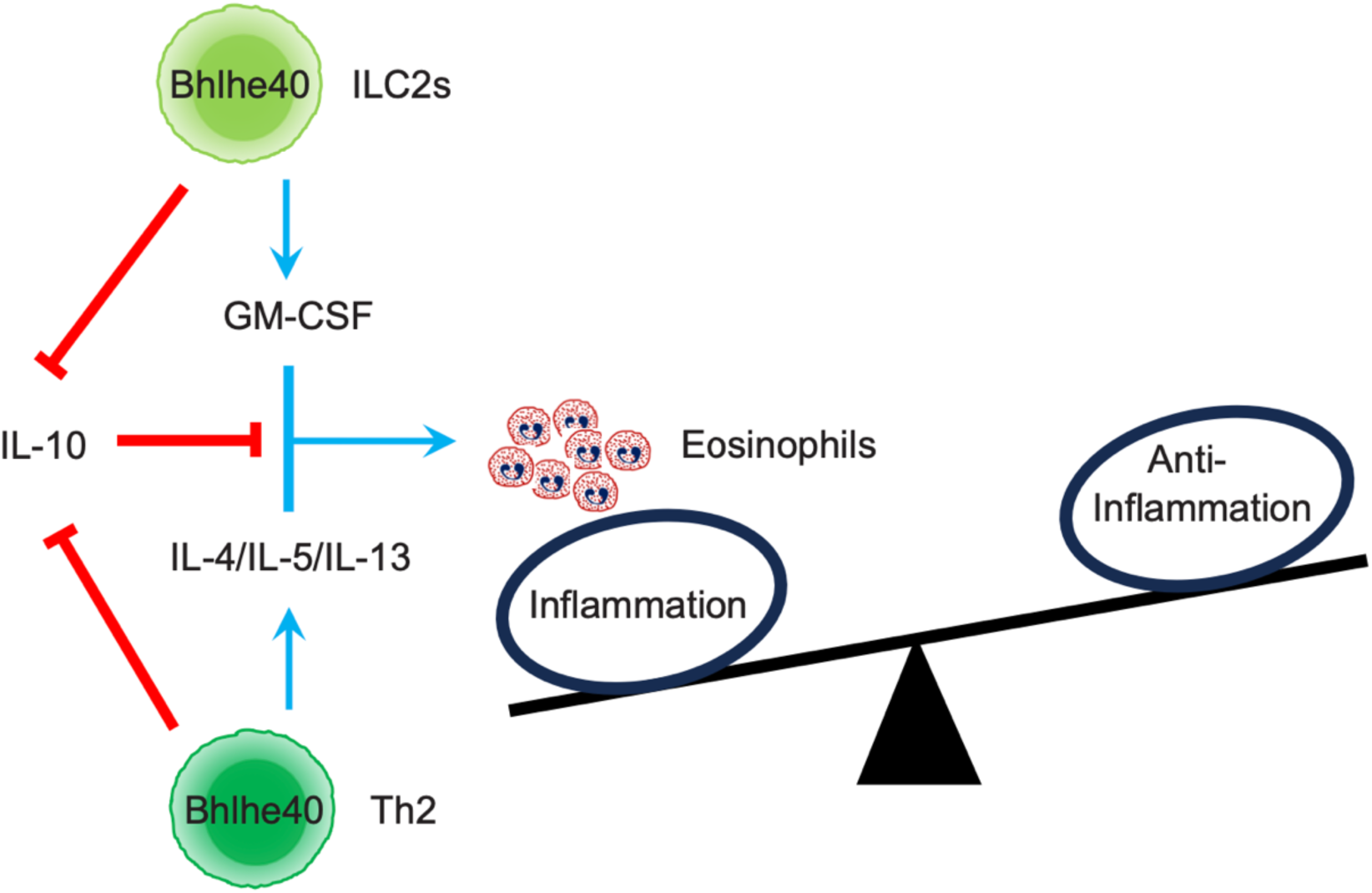
Model for Bhlhe40-mediated gene regulation in ILC2s and Th2 cells during type 2 immune responses. Bhlhe40 is expressed in both ILC2s and Th2 cells. In ILC2s, Bhlhe40 promotes GM-CSF but represses IL-10 production. On the other hand, in Th2 cells, while Bhlhe40 still represses IL-10 production, it also regulates the expression of type 2 cytokines including IL-4, IL-5 and IL-13. Overall, Bhlhe40 promotes eosinophil recruitment by differentially regulating cytokine production by ILC2s and Th2 cells, which results in inflammation in lung.

